# SHIP2 oligomeric states and activity regulate cellular responses to sodium arsenate-induced stress granule dynamics

**DOI:** 10.64898/2026.05.24.727459

**Authors:** Abdulrahman El Sayed, Abdelhalim Azzi

## Abstract

Protein dimerization plays a central role in regulating enzymatic activity, signal transduction, and transcription factor function. Within the PI3K family, different modes of oligomerization have been reported. However, the oligomerization of lipid 5-phosphatases and its functional consequences have not been described. Here, we show for the first time that the lipid 5-phosphatase SHIP2 exists as a homodimer in cells, with its N-terminal region serving as the primary dimerization domain. We further demonstrate that SHIP2 dimerization has no major impact on its catalytic activity but instead profoundly affects its interactome. Interestingly, we identify the stress granule marker G3BP1 as one of the SHIP2 interactors whose association is moderately influenced by SHIP2 oligomerization states. Furthermore, we show that changes in SHIP2 protein levels, enzymatic activity, and oligomeric state alter the cellular response to sodium arsenate-induced stress. In addition, variation in SHIP2 levels and activity affects stress granule size and dynamics. Together, these findings identify SHIP2 oligomerization as a previously unrecognized regulatory mechanism linking phosphoinositide signaling to stress granule biology.

## Introduction

SHIP2 is a multidomain lipid 5-phosphatase that regulates diverse signaling networks. Its domain architecture includes an N-terminal Src Homology 2 (SH2) domain, which mediates interactions with phosphorylated tyrosine residues on signaling proteins such as receptor tyrosine kinases (RTKs); a catalytic inositol 5-phosphatase domain, responsible for hydrolysis of the 5′-phosphate of PI(3,4,5)P3 to generate PI(3,4)P2, thereby modulating PI3K/AKT pathway activity; and a C-terminal Sterile Alpha Motif (SAM) domain that mediates protein-protein and protein-lipid interactions (Elong Edimo et al., 2013; Thomas et al., 2017). The subcellular localization of SHIP2 is highly dynamic, enabling it to participate in diverse cellular processes. SHIP2 regulates cellular responses to extracellular stimuli through both enzymatic and adaptor functions. For example, recruitment of SHIP2 to the plasma membrane during receptor activation promotes local production of PI (3,4) P2, which is required for clathrin-mediated endocytosis (Chan Wah Hak et al., 2018; Nakatsu et al., 2010). In parallel, its lipid phosphatase activity suppresses PI3K/AKT signaling by hydrolyzing PI (3,4,5) P3, thereby modulating the PI (3,4,5) P3/ PI (3,4) P2 balance and limiting AKT activation. This regulatory mechanism is considered essential for maintaining cellular homeostasis and preventing hyperactivation of growth and survival signals (Kerr, 2011). In cancer cells, SHIP2 promotes invadopodia formation, facilitating matrix degradation and metastatic dissemination. It also regulates the turnover of growth factor receptors, including EGFR and FGFR, through interactions with the ubiquitin ligase CBL (Fafilek et al., 2018; Prasad, 2009; Vandenbroere et al., 2003). Beyond its cytoplasmic functions, a phosphorylated form of SHIP2 has been detected at nuclear speckles, dynamic, membrane-less compartments enriched in RNA-processing factors, suggesting a potential role in RNA metabolism and transcriptional regulation (Elong Edimo et al., 2011). However, the contribution of SHIP2 to RNA-associated mechanisms remains largely unexplored. Biomolecular condensates are membrane-less cellular assemblies of proteins and nucleic acids, collectively referred to as ribonucleoprotein (RNP) granules (Millar et al., 2023). These structures form via liquid-liquid phase separation (LLPS), in which dispersed, protein-associated RNA molecules transition from a soluble state to a condensed phase with liquid-like properties. Nuclear speckles and cytoplasmic RNP granules share common organizational principles, suggesting that SHIP2 may function within phase-separated RNA-protein assemblies. Stress granules (SGs) are a prominent class of RNP granules that assemble in the cytoplasm of eukaryotic cells in response to diverse stimuli, including oxidative stress, heat shock, and proteotoxic stress. A central molecular event in SG formation is the stress-induced phosphorylation of eukaryotic initiation factor 2α (eIF2α), which inhibits translation initiation and triggers the cytoplasmic accumulation of untranslated mRNPs (Buchan and Parker, 2009). SG formation and disassembly are tightly regulated processes that influence mRNA fate, translational control, and cellular health. Key RNA-binding proteins (RBPs), including TIA-1, G3BP1, and G3BP2, have been identified as essential nucleators of SG assembly, acting alongside translation initiation factors such as eIF3 and eIF4A and the associated RBP CAPRIN1 (Millar et al., 2023). These proteins undergo regulated conformational changes that govern their interactions, assembly behavior, and subcellular localization. In particular, protein oligomerization plays a central role in modulating interaction specificity and driving condensate formation. For example, G3BP1 dimerizes through its NTF2-like (NTF2L) domain, and this self-association is essential for SG nucleation and RNA metabolism (Sidibé et al., 2021; Vognsen et al., 2013). The importance of dimerization as a regulatory mechanism extends beyond SG biology. Indeed, dimerization of the p110α and p85α subunits of class I PI3K stabilizes its catalytic activity (Cheung et al., 2015), and homodimerization of the phosphatase PTEN is required for its full lipid phosphatase activity (Heinrich et al., 2015). However, whether oligomerization similarly regulates the activity and function of phosphoinositide 5-phosphatases remains unknown. In this study, we demonstrate that SHIP2 forms homodimers through its N-terminal domain in cells. Proteomic analysis further shows that the SHIP2 interactome is markedly influenced by its oligomeric state. Interestingly, the central stress granule regulator G3BP1 emerges among the top SHIP2 interactors. We further show that modulation of SHIP2 oligomerization and phosphatase activity significantly impacts stress granule assembly and dynamics. Together, these findings identify SHIP2 oligomerization as a previously unrecognized regulatory mechanism linking phosphoinositide signaling to stress granule biology.

## Results

### SHIP2 exists in a dimeric complex in vivo via its N-terminal domain

SHIP2 is a large multidomain protein whose full-length three-dimensional structure has not yet been resolved by X-ray crystallography, largely owing to its extensive intrinsically disordered regions. Structural predictions generated using AlphaFold2 (Jumper et al., 2021) provide insight into the overall domain organization and highlight the challenges associated with resolving its complete conformation (**Figure 1 A-B**). The predicted model confirms the presence of a well-defined SH2 domain, a RhoA-binding domain, and a pleckstrin homology-related (PH) domain within the N-terminal region, followed by the phosphatase core and a C-terminal sterile alpha motif (SAM) domain (Thomas et al., 2017). As illustrated in **Figure 1 B**, SHIP2 harbors extensive disordered regions, suggesting a high degree of conformational flexibility that may facilitate interactions with multiple binding partners and enable context-dependent regulation of SHIP2 activity.

**Figure 1:**
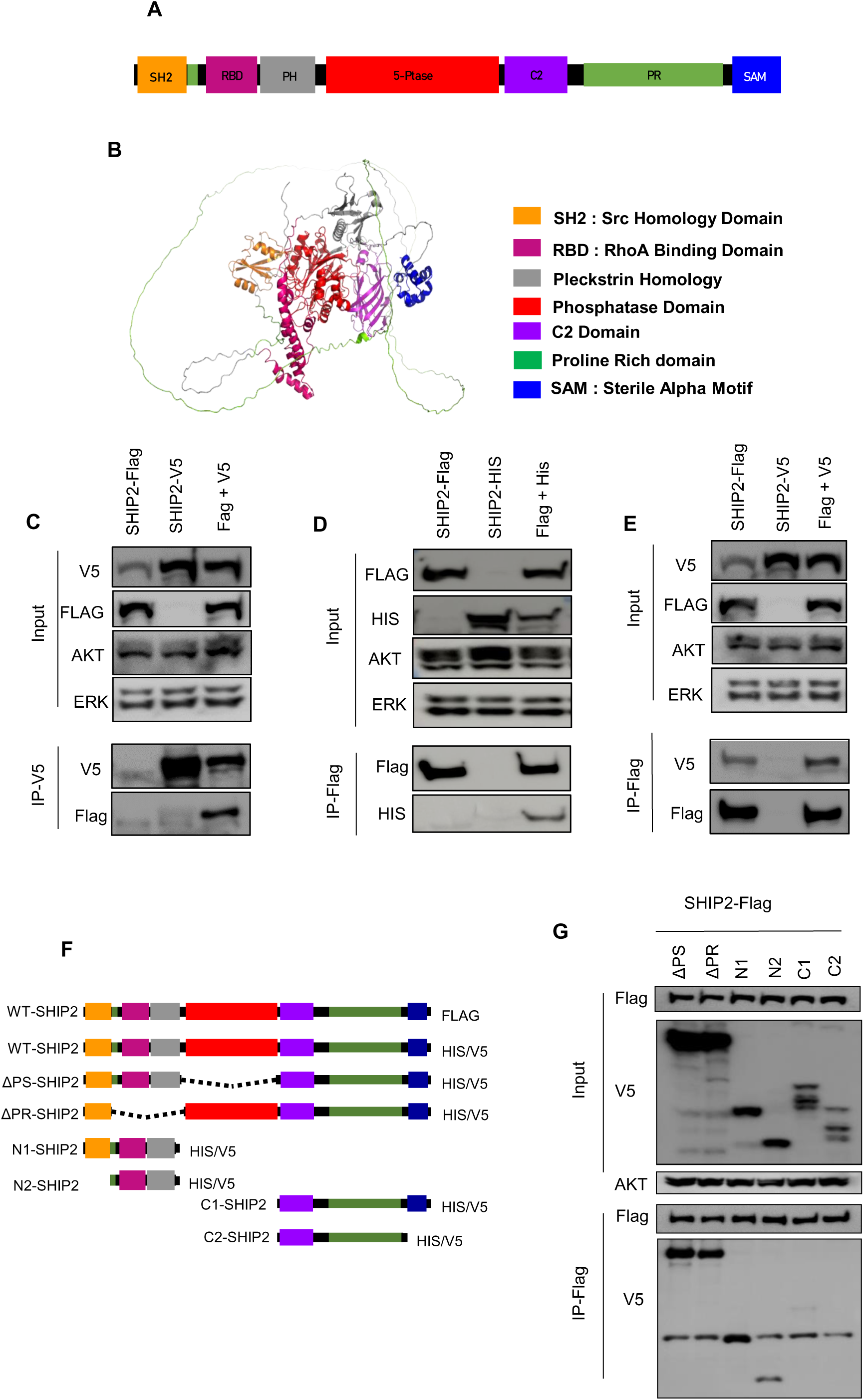
SHIP2 forms homodimers through its N-terminal domain. (**A–B**) Schematic representation and AlphaFold2-predicted structural model of SHIP2 and its domain organization. The SH2 domain is shown in yellow, proline-rich (PR) domains in green, pleckstrin homology (PH) domain in grey, RhoA-binding domain (RBD) in purple, phosphatase domain in red, and sterile alpha motif (SAM) in blue. (**C–E**) SHIP2 forms homodimers in cells. Flag- or V5/His-tagged SHIP2 constructs were co-expressed in HEK293T cells. Forty-eight hours post-transfection, cells were lysed and immunoprecipitated with antibodies against the Flag (D), V5 (C), or His (E) epitope tags and analyzed by western blotting. AKT and ERK were used as loading controls. Input lanes represent 10% of the total lysate used for immunoprecipitation. Representative of 3 biological replicates. (**F**) Schematic representation of the SHIP2 deletion constructs used for domain mapping. ΔPS: phosphatase domain deletion (Δ419–739 aa); ΔPR: proline-rich domain deletion (Δ123–410 aa); N1: N-terminal fragment (Δ419–1258 aa); N2: N-terminal fragment lacking the SH2 domain (Δ1–122, 419–1258 aa); C1: C-terminal fragment (Δ1–739 aa); C2: C-terminal fragment lacking the SAM domain (Δ1–739, 1190–1258 aa). (**G**) Domain-mapping of the SHIP2 dimerization interface. The indicated His/V5-tagged SHIP2 deletion constructs were co-expressed in HEK293T cells with Flag-tagged full-length SHIP2, immunoprecipitated using an anti-V5 antibody, and analyzed by western blotting. Input is the total cell lysate used for immunoprecipitation. Representative of 3 biological replicates.

To determine whether SHIP2 undergoes homodimerization in cells, Flag-tagged SHIP2 and His/V5-tagged SHIP2 were transiently co-expressed in HEK293T cells, and co-immunoprecipitation experiments were performed using antibodies against the V5 (**Figure 1 C**), Flag (**Figure 1 D**), and His (**Figure 1 E**) epitope tags. In each condition, the co-expressed SHIP2 variant was co-precipitated, demonstrating that SHIP2 can self-associate and form homodimers in cells. To map the domain(s) responsible for homodimerization, a panel of His/V5-tagged SHIP2 deletion plasmid constructs were used. The deletions include the phosphatase domain (ΔPS), the proline-rich domain (ΔPR), as well as isolated N-terminal (N1, N2) and C-terminal (C1, C2) fragments (**Figure 1 F**). Each construct was co-expressed with full-length Flag-tagged SHIP2, and co-immunoprecipitations were performed using an anti-Flag antibody followed by immunodetection of the V5 tag. As shown in Figure 1G, only constructs retaining the N-terminal region yielded a detectable co-precipitated band, whereas C-terminal fragments did not associate with full-length SHIP2. Collectively, these data establish that SHIP2 forms homodimers in vivo and that the N-terminal domain is necessary for this self-association.

### SHIP2 catalytic activity and interactome depend on its oligomeric state

SHIP2 negatively regulates PI (3,4,5) P3 levels through its phosphatase activity, while also functioning as a crucial adaptor protein for sustained ERK signaling (Fafilek et al., 2018). To investigate whether the oligomeric state of SHIP2 influences its activity and interaction network, we first generated a dimer-deficient SHIP2 mutant (referred to hereafter as ΔN-SHIP2) by deleting the N-terminal domain (aa 21–427) through site-directed mutagenesis, while preserving the integrity of the remaining protein (**Figure 2 A**). Co-immunoprecipitation of Flag-tagged ΔN-SHIP2 co-expressed with its His-tagged wild-type counterpart confirmed a complete loss of self-association for the mutant, independently validating the results obtained above showing that the N-terminal deletion selectively abrogates homodimerization (**Figure 2 B**).

**Figure 2:**
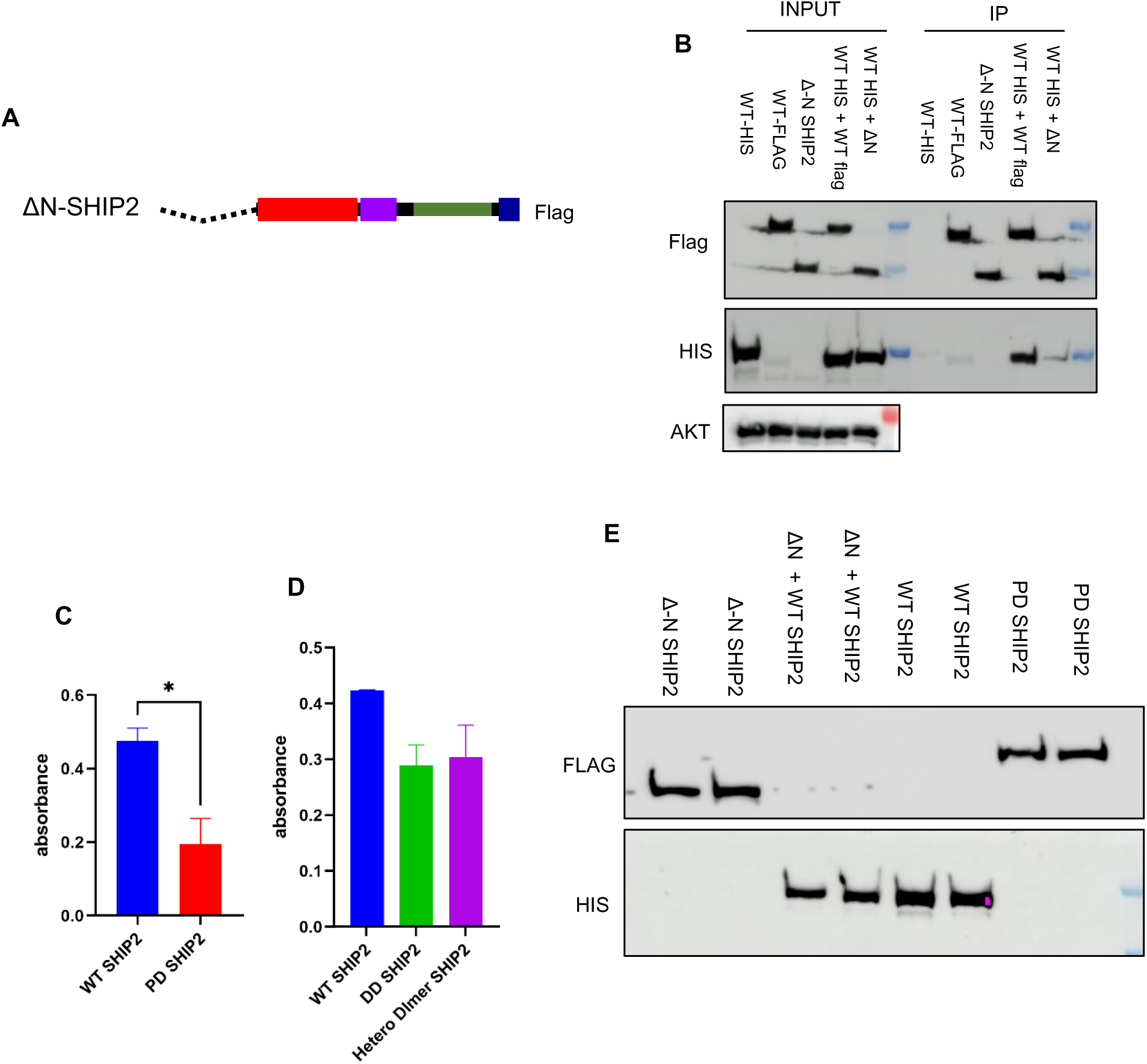
Disruption of SHIP2 homodimerization modestly reduces its phosphatase activity. (**A**) Schematic representation of the Flag-tagged ΔN-SHIP2 deletion mutant (Δ21–427 aa). (**B**) ΔN-SHIP2 exhibits abrogated homodimerization. Wild-type Flag-tagged SHIP2, wild-type His-tagged SHIP2, and Flag-tagged ΔN-SHIP2 were co-expressed in HEK293T cells. Forty-eight hours post-transfection, cells were immunoprecipitated using an anti-Flag antibody and analyzed by western blotting. Total AKT was used as a loading control. (**C–D**) The oligomeric state of SHIP2 moderately influences its phosphatase activity. Wild-type, phosphatase-dead, ΔN-SHIP2, and a combination of ΔN-SHIP2 and wild-type SHIP2 were transiently expressed in HEK293T cells. Twenty-four hours post-transfection, each form was immunoprecipitated using the corresponding epitope-tag antibody; bead-bound complexes were incubated with PI (3,4,5) P3, and released inorganic phosphate was quantified using a malachite green assay. Beads were subsequently resuspended in Laemmli buffer, and immunoprecipitated proteins were resolved by SDS-PAGE to confirm expression levels and precipitation (**E**).

We next examined whether oligomerization influences SHIP2 catalytic activity. Phosphatase activity was assessed using an on-bead assay with PI (3,4,5) P3 as a substrate and malachite green-based detection of released inorganic phosphate. Wild-type and phosphatase-dead (PD) SHIP2 were immunoprecipitated via their respective epitope tags and incubated with substrate; PD-SHIP2 served as the negative control. This approach showed that the catalytic activity of PD-SHIP2 was reduced to approximately half that of wild-type SHIP2 (**Figure 2 C**), establishing the assay dynamic range. When ΔN-SHIP2 or the combination of wild-type and ΔN-SHIP2 (heterodimeric state) were assessed using the same assay, a modest but reproducible reduction in phosphatase activity was observed relative to wild-type SHIP2 alone (**Figure 2 D**). These findings suggest that disruption of SHIP2 homodimerization partially impairs catalytic efficiency. This effect is consistent with an allosteric or interdomain contribution of the N-terminal region to the optimal orientation or activity of the phosphatase domain, rather than a direct catalytic role. Importantly, the partial enzymatic activity retained by both ΔN-SHIP2 and the heterodimeric form indicates that the phosphatase domain remains intrinsically functional in the absence of the dimerization interface.

To determine how SHIP2 oligomerization influences its interaction landscape, we performed affinity purification coupled to mass spectrometry (AP–MS) in HEK293T cells under the following conditions: wild-type His-tagged SHIP2 co-immunoprecipitated using an anti-Flag antibody, wild-type His-tagged SHIP2 co-expressed with Flag-tagged ΔN-SHIP2 and immunoprecipitated using an anti-His antibody, or Flag-tagged ΔN-SHIP2 alone immunoprecipitated using an anti-Flag antibody (**Figure 3 A-C**). This approach revealed that the SHIP2 interactome varies with its oligomeric state. Indeed, both the number and functional profiles of the proteins associated with each SHIP2 oligomeric state differ, especially in the dimer-deficient SHIP2 (**Figure 3 A-C and supplementary table 1**). Gene Ontology (GO) analysis of the interactome associated with dimeric wild-type SHIP2 revealed that top represented pathway was related to ubiquitin protein ligase binding, including proteins involved in protein folding, chaperone activity, and cytoskeletal regulation (**Figure 3 D, supplementary table 2**). In contrast, the interactome of ΔN-SHIP2 showed a pronounced shift toward RNA-binding and RNA-metabolic functions (**Figure 3 F**). Notably, co-expression of wild-type SHIP2 with ΔN-SHIP2, followed by immunoprecipitation of wild-type SHIP2, yielded a molecular function profile that closely resembled that of ΔN-SHIP2 alone (**Figure 3 E, supplementary table 2**), suggesting that the presence of the monomeric form exerts a dominant influence on the interaction landscape. Interestingly, across all conditions, a subset of proteins interacting with SHIP2 was consistently detected, albeit at varying enrichment levels; these shared interactors were predominantly associated with stress granule formation and RNA metabolism, most notably G3BP1 (**Supplementary table 1**). This result suggests that SHIP2 may participate in a core interaction module linked to stress granule dynamics and RNA metabolism that is preserved across oligomeric states, while the broader interactome is selectively altered depending on its oligomerization status. Together, these findings indicate that SHIP2 oligomerization governs both its catalytic efficiency and its engagement with distinct protein networks. The dynamic balance between dimeric and monomeric SHIP2 may serve as a molecular switch, redirecting its cellular activities depending on the nature of the stress stimulus.

**Figure 3:**
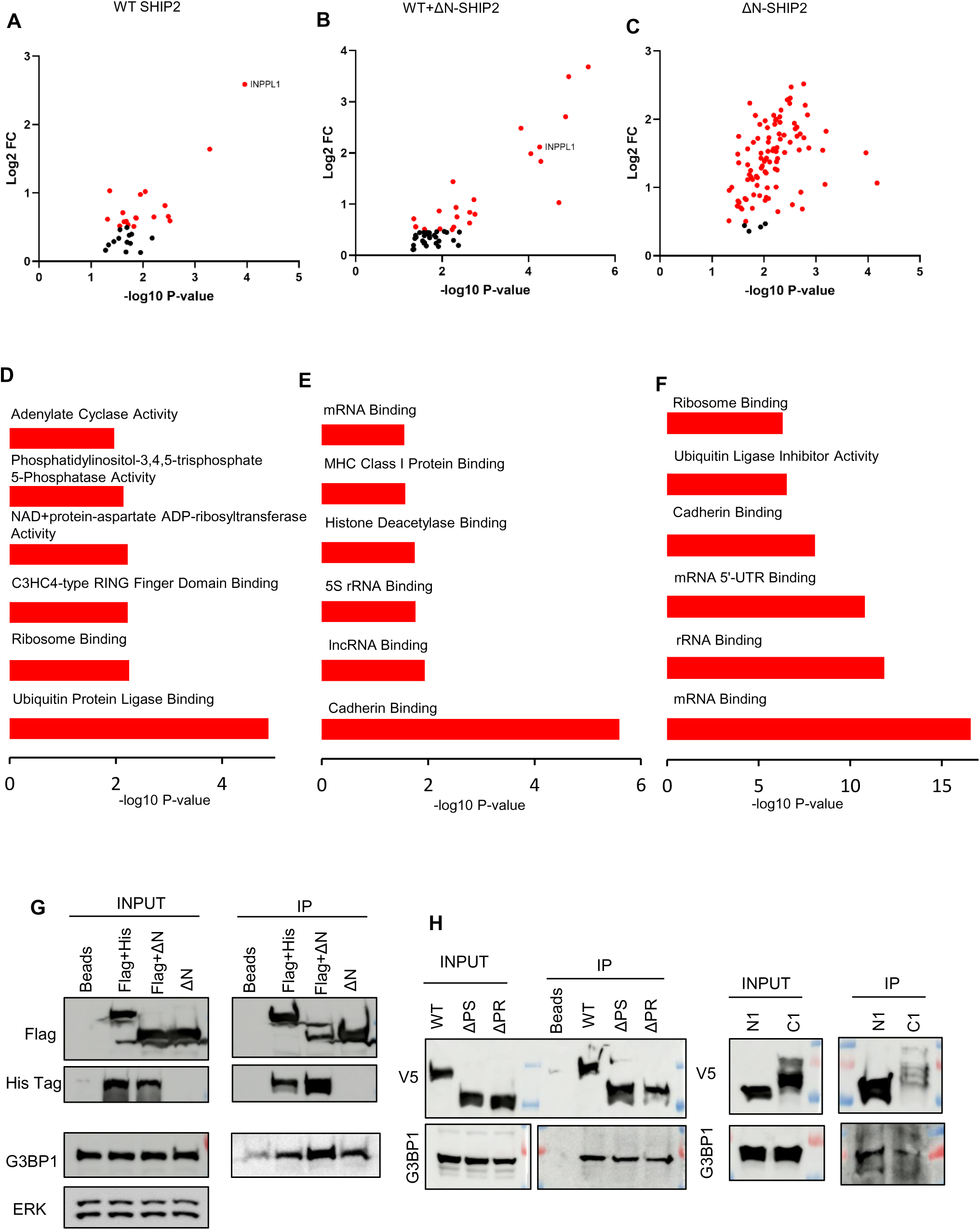
The SHIP2 interactome is dependent on its oligomeric state. (**A–C**) Volcano plots of proteins identified by AP-MS analysis of immunoprecipitated SHIP2 complexes. The X-axis represents −log10 p-value; the Y-axis represents log2 fold change relative to the IgG control. Enrichment threshold was defined at 0.5 and the significance threshold was set to >1.3 -log10 P-value, N = 3 biological replicates/condition (**A**) Wild-type His-tagged SHIP2 and Flag-tagged SHIP2 co-expressed and immunoprecipitated using an anti-His antibody. (**B**) Wild-type His-tagged SHIP2 co-expressed with Flag-tagged ΔN-SHIP2, immunoprecipitated using an anti-Flag antibody. (**C**) Flag-tagged ΔN-SHIP2 was expressed alone and immunoprecipitated using an anti-Flag antibody. (**D–F**) Gene Ontology (GO) analysis of molecular functions enriched in the interactomes of wild-type SHIP2 (**D**), wild-type + ΔN-SHIP2 co-expression (**E**), and ΔN-SHIP2 alone (**F**). The Y-axis lists molecular function terms; the X-axis shows −log10 enrichment p-values. (**G**) Independent validation of the SHIP2 interaction with the stress granule marker G3BP1. Wild-type Flag-tagged SHIP2, His-tagged SHIP2, and Flag-tagged ΔN-SHIP2 were co-expressed in HEK293T cells and immunoprecipitated using an anti-Flag antibody. Blots were probed with the indicated antibodies. Total ERK was used as a loading control. Representative of 3 biological replicates. (**H**) Mapping the G3BP1-binding site on SHIP2. The indicated Flag-tagged SHIP2 constructs were expressed in HEK293T cells, immunoprecipitated using an anti-Flag antibody, and probed for G3BP1 by western blotting. Representative of 3 biological replicates.

### SHIP2 interacts with stress granule markers

To independently validate the SHIP2 interactions with G3BP1 identified by AP–MS, endogenous SHIP2 was immunoprecipitated from non-transfected HEK293T cells using a SHIP2-specific antibody, and co-precipitated proteins were analyzed by mass spectrometry. As shown in supplementary table 3, this approach validated the interaction of endogenous SHIP2 with CBL, a ubiquitin ligase previously reported to associate with SHIP2 (Vandenbroere et al., 2003), and, importantly, confirmed the interaction of SHIP2 with G3BP1. These findings establish that SHIP2 associates with stress granule components under endogenous expression conditions and is not dependent on exogenous overexpression.

Furthermore, co-immunoprecipitation experiments using an anti-Flag antibody in cells co-expressing Flag-tagged wild-type SHIP2, His-tagged wild-type SHIP2, and Flag-tagged ΔN-SHIP2 further validated these interactions. Indeed, western blot analysis confirmed the association of SHIP2 with G3BP1, with interaction levels varying between the wild-type and dimer-deficient forms (**Figure 3 G**). Consistent with our observations, SHIP2 has previously been reported to localize to perinuclear speckles (Elong Edimo et al., 2011), a compartment associated with RNA metabolism, further supporting a role for SHIP2 in RNA granule biology.

To characterize the domain of SHIP2 responsible for the interaction with G3BP1, a panel of Flag-tagged SHIP2 plasmid constructs were used. This includes the wild-type SHIP2, the phosphatase domain deletion (ΔPS), the proline-rich domain deletion (ΔPR), and isolated N-terminal and C-terminal fragments, which were used in co-immunoprecipitation experiments followed by G3BP1 immunodetection. The interaction with G3BP1 was retained in constructs harboring the N-terminal domain of SHIP2 (**Figure 3 H**). However, our interactome experiments (Supplementary Table 1) reveal that the SHIP2–G3BP1 interaction is retained even upon deletion of the N-terminal domain. These data suggest that, while G3BP1 preferentially binds SHIP2 through the N-terminal domain, a residual interaction can still occur via an alternative domain, particularly given that removal of the N-terminal domain may alter the conformational landscape of the remaining SHIP2 domains.

### SHIP2 colocalizes with stress granule components

To validate the physical association of SHIP2 with stress granule components in living cells, Flag-tagged wild-type SHIP2 was expressed in HEK293T cells. Sodium arsenate, a well-established and potent inducer of stress granules (Bernstam and Nriagu, 2000), was used to induce stress granule formation. Cells were analyzed by confocal immunofluorescence microscopy using antibodies against Flag and G3BP1 as a canonical SG marker. Under basal, unstressed conditions, a partial colocalization between SHIP2 (red) and G3BP1 (green) was observed (**Figure 4 A**), consistent with a low-level, constitutive association. Following 15 minutes of sodium arsenate treatment, the degree of colocalization markedly increased (**Figure 4 B**), indicating that stress promotes the recruitment of SHIP2 to nascent G3BP1-containing foci. Notably, after 30 minutes of treatment, at a time point when mature, phase-separated stress granules are established, SHIP2 was no longer detected within the G3BP1-positive granule cores but instead concentrated at the granule periphery (**Figure 4 C**). Signal intensity profiles confirmed this redistribution (**Figure 4 D–F**). These data suggest that SHIP2 participates in the early stages of SG assembly and subsequently adopts a peripheral localization at mature condensates, consistent with a role in regulating SG dynamics at the condensate-cytoplasm interface rather than as a stable core component.

**Figure 4:**
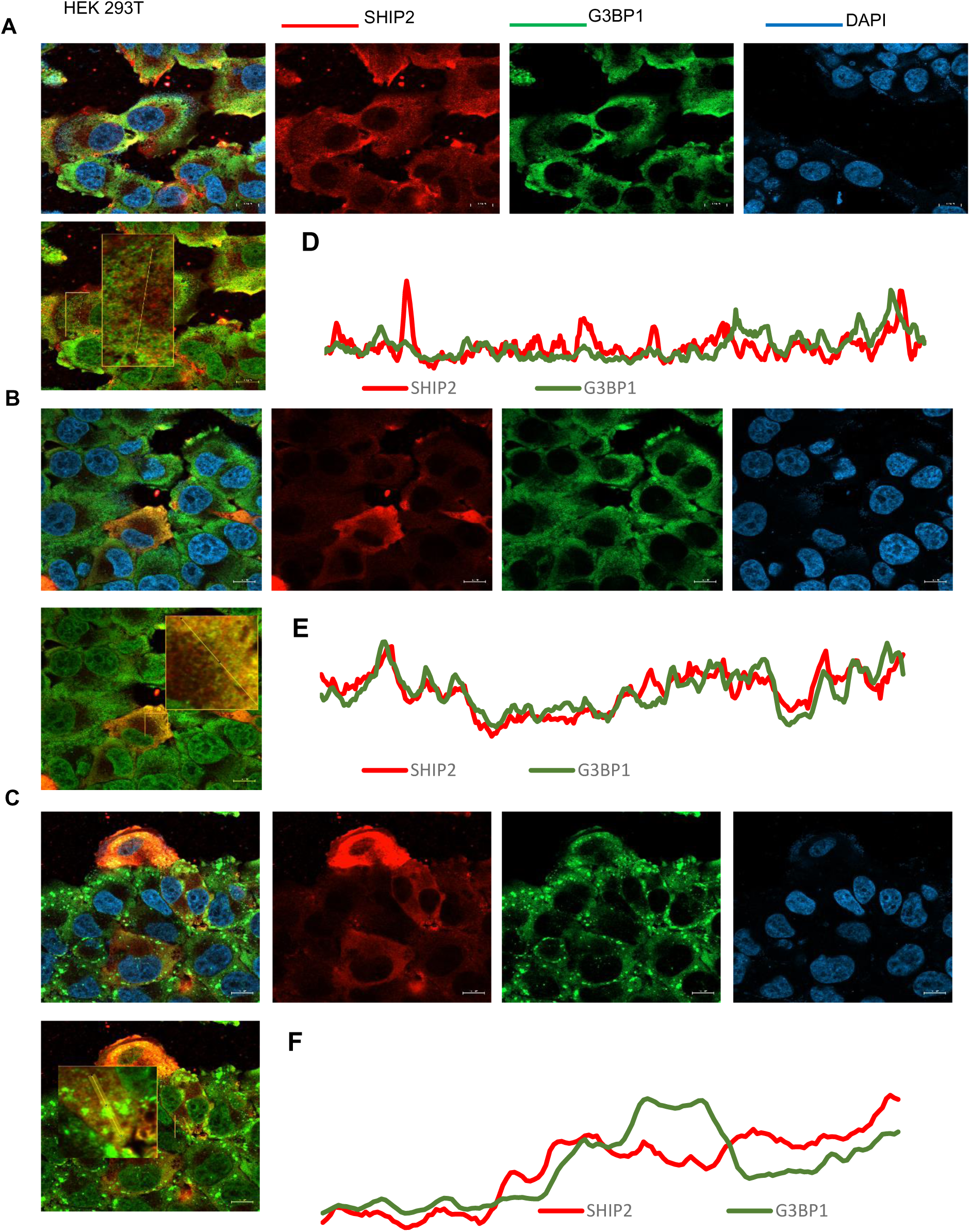
SHIP2 colocalizes with G3BP1 during stress and localizes to the periphery of mature stress granules. (**A–C**) HEK293T cells transiently expressing His-tagged wild-type SHIP2 were left untreated (**A**) or treated with 500 μM sodium arsenate for 15 minutes (**B**) or 30 minutes (**C**). Cells were fixed and stained for SHIP2 (red), G3BP1 (green), and DAPI (blue). Images were acquired by confocal microscopy. (**D–F**) Signal intensity profiles of SHIP2 (red) and G3BP1 (green) fluorescence across representative cells from the untreated (**D**), 15-minute (**E**), and 30-minute (**F**) arsenate-treated conditions.

### Changes in SHIP2 levels, activity, and oligomeric state alter cellular response to sodium arsenate

Studies have shown that sodium-arsenate mediated stress granule formation exerts a global impact on protein synthesis and mRNA translation (Buchan and Parker, 2009; Kedersha et al., 2016). To determine how different SHIP2 oligomeric states influence the broader cellular proteome in response to stress, HEK293T cells transiently expressing an empty vector, wild-type SHIP2, phosphatase-dead SHIP2, wild-type + ΔN-SHIP2 (heterodimeric condition), or ΔN-SHIP2 alone were treated with 500 μM sodium arsenate for two hours. Total cell lysates were then subjected to quantitative mass spectrometry (**Figure 5 A-C, G-H**). This analysis revealed that changes in SHIP2 levels and its oligomeric state exert a global effect on arsenate-mediated stress. Indeed, the number of significantly altered proteins and the enriched molecular functions differed substantially across conditions (**Figure 5, supplementary table 4**). This analysis revealed that in control cells expressing an empty vector, arsenate treatment upregulated DNA oriented processes and DNA-template transcription, while downregulating protein catabolism and metabolic related processes, reflecting the canonical cellular response to oxidative stress. In contrast, expression of wild-type SHIP2 shifted this response toward downregulation of cellular development, mTORC1 signaling, and macroautophagy, with concurrent upregulation of proteins involved in mitochondrial respiration and DNA replication. Interestingly, phosphatase-dead SHIP2 expression was associated with downregulation of mTORC1 signaling and splicing, while simultaneously upregulating translation and protein biosynthetic processes (**Figure 5 A-F**). Together, these data suggest that the lipid phosphatase activity and the adaptor function of SHIP2 differentially modulate the cellular response to arsenate-induced stress. We next examined the impact of SHIP2 oligomeric state. As indicated in (**Figure 5 H, J**), the dimer-deficient form of SHIP2 (ΔN-SHIP2) uniquely downregulated RNA-related functions while upregulating DNA-related processes and splicing pathways. In contrast, the heterodimeric state was primarily characterized by downregulation of endocytosis and intracellular organization, alongside upregulation of endoplasmic reticulum stress-related processes (**Figure 5 G-I**). Taken together, these findings demonstrate that both the enzymatic activity and oligomeric state of SHIP2 differentially shape the global proteomic response to arsenate-induced stress, highlighting its role as a multifunctional regulator that integrates lipid signaling, RNA metabolism, and stress-responsive translational control.

**Figure 5:**
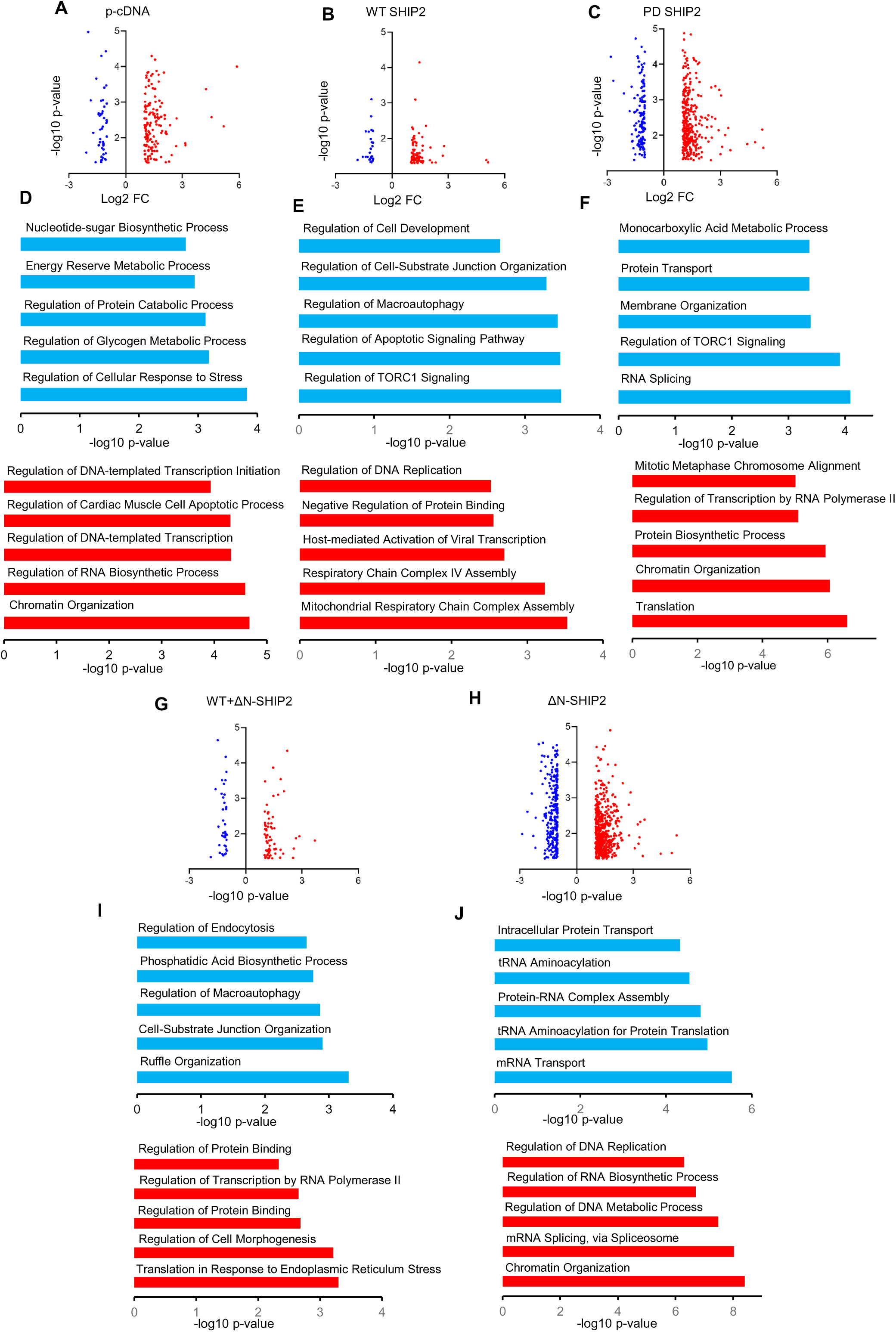
Changes in SHIP2 levels and oligomeric state remodel the stress-regulated proteome. (**A–C, G-H**) Volcano plots of differentially expressed proteins identified by quantitative mass spectrometry analysis of total lysates from HEK293T cells transiently expressing pcDNA empty vector **(A)**, wild-type SHIP2 **(B)**, phosphatase-dead SHIP2 **(C)**, wild-type + ΔN-SHIP2 **(G)**, or ΔN-SHIP2 alone **(H)**, following treatment with 500 μM sodium arsenate. The Y-axis represents −log10 p-value; the X-axis represents log2 fold change relative to a non-treated control. 3 biological replicates were analyzed condition. (**D-F, I–J**) Gene Ontology analysis of significantly up- (red) and downregulated (blue) molecular functions in pcDNA control **(D)**, wild-type SHIP2 **(E)**, phosphatase-dead SHIP2 **(F)**, wild-type + ΔN-SHIP2 **(I)**, and ΔN-SHIP2 **(J)** expressing cells under arsenate-induced stress. The X axis represents -log10 P-value and the Y axis indicates Molecular Function terms.

### Changes in SHIP2 levels and activity alter stress granule size and dynamics

Having established that SHIP2 associates with stress granules and that modulation of its expression and catalytic activity influences the cellular response to sodium arsenate, we next sought to quantify its impact on SG size and assembly dynamics. MDA-MB-231 cells stably overexpressing wild-type or phosphatase-dead SHIP2, together with parental MDA-MB-231 cells as controls (**Figure 6 A–C**), were treated with 500 μM sodium arsenate for 1 hour, fixed, and immunostained for G3BP1 to visualize stress granules. A minimum of 35 cells per condition were analyzed. The size distribution of SGs was broadly comparable across conditions for granules smaller than 3 μm in diameter (**Supplementary Figure 1 A–C**); however, cells overexpressing wild-type SHIP2 displayed a greater number and larger proportion of SGs exceeding 3 μm relative to both parental and phosphatase-dead SHIP2-expressing cells (**Figure 6 D–F**). This relationship was inverted upon SHIP2 knockdown: cells expressing shRNA targeting SHIP2 showed a significantly lower number and reduced size of SGs >3 μm compared to non-targeting shControl-expressing cells (**Figure 6 G–L**), whereas the distribution of smaller granules remained largely unchanged (**Supplementary Figure 1 D–F**).

**Figure 6:**
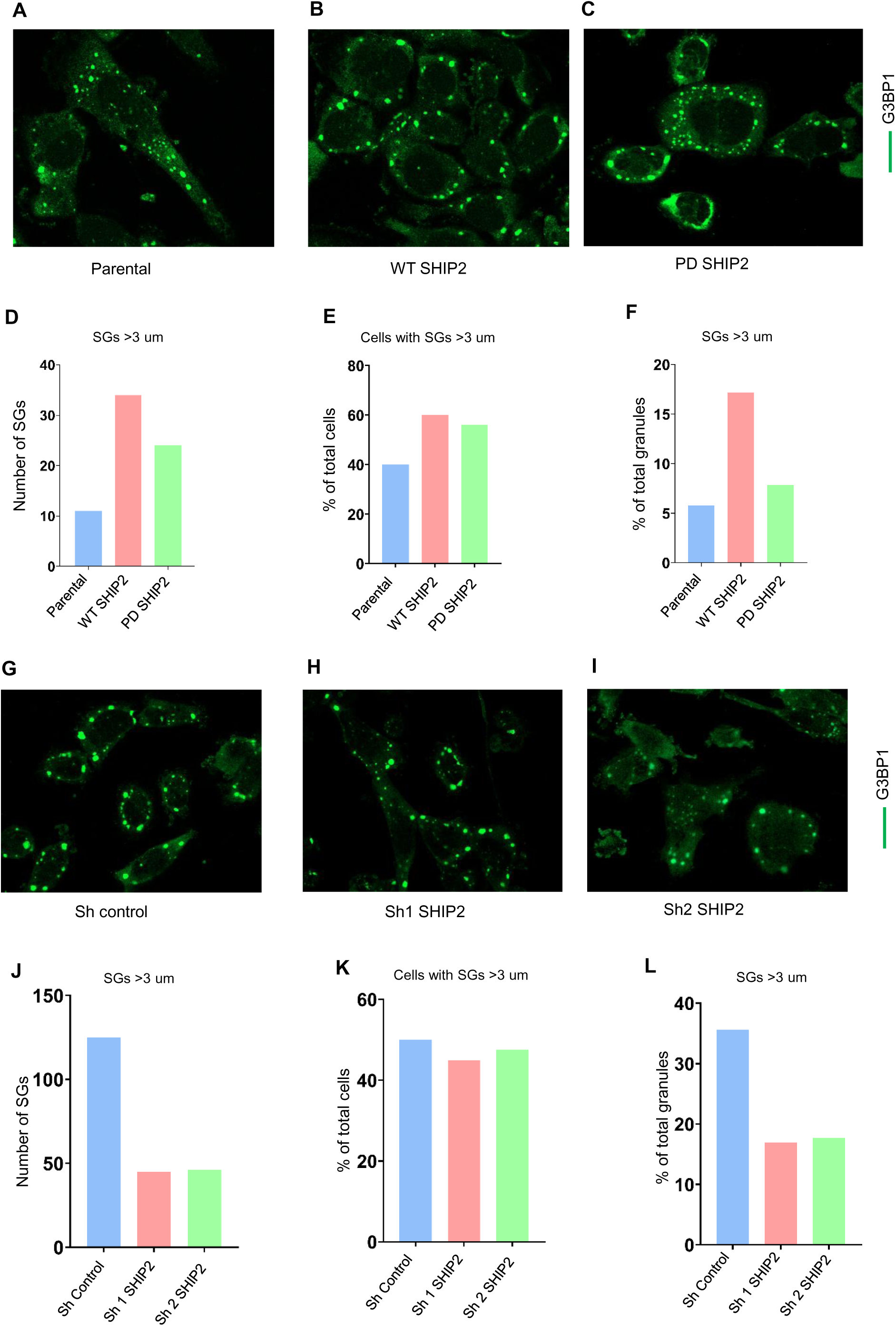
SHIP2 expression and catalytic activity alter stress granule size. **(A–C)** confocal images of MDA-MB-231 under arsenate stress. Parental cells **(A)**, and cells stably overexpressing wild-type SHIP2 **(B)** or phosphatase-dead SHIP2 **(C)**, **(G-I)** cells expressing Sh control **(G)**, and SHIP2 knockdown using Sh1 **(H)** and Sh2 **(I)**, were treated with 500 μM sodium arsenate for one hour, washed with PBS, and fixed with 4% paraformaldehyde. Cells were stained for G3BP1 (green). Scale bar, 10 μm. **(D–F, J-L)** Quantification of stress granules. Bar graphs representing the number of SGs >3 um in corresponding cell lines **(D, J)**, Percentage of cells exhibiting SGs >3 um **(E, K)**, and Percentage of SGs >3 in each cell line. **(F, L)** 40 cells per condition were analyzed.

We next examined the effect of SHIP2 expression and catalytic activity on stress granule dynamics. Following 1 hour of sodium arsenate treatment, the proportion of G3BP1-positive cells was markedly lower in cells stably overexpressing wild-type SHIP2 relative to parental controls (∼20% reduction), with phosphatase-dead SHIP2-expressing cells exhibiting an intermediate reduction (∼10%) (**Figure 7 A, E, I**). Prolonged arsenate exposure further differentiated these responses: wild-type SHIP2-overexpressing cells retained larger and more numerous stress granules after 3 hours of treatment compared to phosphatase-dead SHIP2 cells and parental controls (**Supplementary Figure 2 A–J**), indicating that both the catalytic and adaptor functions of SHIP2 contribute to sustained SG maintenance under persistent stress.

**Figure 7:**
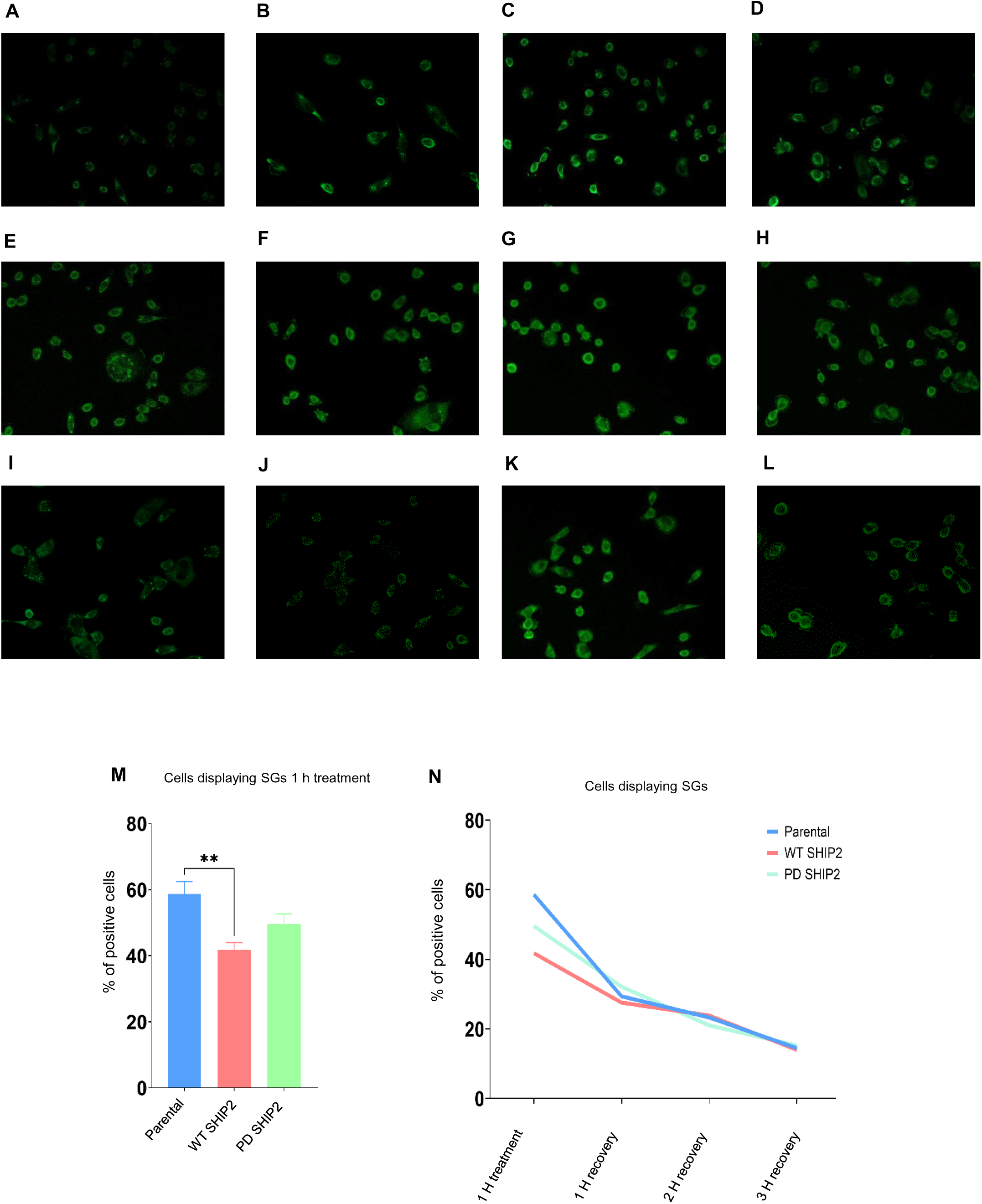
SHIP2 expression and catalytic activity alter cellular response to stress granule formation. (A–L) Confocal images of MDA-MB-231 cells immunostained for G3BP1 (green). Parental cells after 1 hour of arsenate treatment **(A)** or 1 hour **(B)**, 2 hours **(C)**, and 3 hours **(D)** of recovery; cells stably overexpressing wild-type SHIP2 after 1 hour of treatment **(E)** or 1 hour **(F)**, 2 hours **(G)**, and 3 hours **(H)** of recovery; and cells stably overexpressing phosphatase-dead SHIP2 after 1 hour of treatment **(I)** or 1 hour **(J)**, 2 hours **(K)**, and 3 hours **(L)** of recovery. **(M)** Quantification of stress granule formation in response to arsenate treatment. The percentage of cells positive for stress granules is shown for each cell line. N = 190 cells per condition. Y axis represents the percentage of cells positive for SGs and the X axis indicates the cells line **(N)** Quantification of stress granule clearance kinetics. The percentage of cells positive for stress granules is shown following 1 hour of arsenate treatment and at 1, 2, and 3 hours of recovery for each cell line. the Y axis represents the percentage of cells, and the X axis displays the condition.

To assess SG clearance kinetics, cells were allowed to recover following a 1-hour arsenate treatment. At 1 and 2 hours post-washout, stress granule clearance proceeded at comparable rates across all cell lines. By 3 hours of recovery, the proportion of G3BP1-positive cells had converged to similar levels across all conditions (**Figure 7 N**), suggesting that SHIP2 does not substantially influence the kinetics of SG disassembly. Taken together, these findings indicate that SHIP2 expression levels and phosphatase activity preferentially regulate the early stress response and the assembly of large stress granules, while playing a limited role in their subsequent clearance.

## Discussion

Protein dimerization is a crucial biological process involving specific protein domains. This process directly regulates enzymatic activity, signal transduction, and transcription factor activation. Different modes of dimerization or oligomerization within the PI3K family have been reported (Harpur et al., 1999; Pérez-García et al., 2014). Moreover, studies have shown that dimerization is essential for the activity of the lipid phosphatase PTEN (Papa et al., 2014). Despite the established importance of dimerization in various families of lipid phosphatases, little is known about the dimerization of lipid 5-phosphatases and the functional consequences of such interactions. In the current study, we show for the first time that lipid 5-phosphatase SHIP2 exists as a homodimer in cells, with its N-terminal domain serving as the primary interface mediating self-association. This finding adds a fundamentally new layer to the regulatory landscape of SHIP2, which to date has been largely described in terms of its catalytic phosphatase activity and its roles as an adaptor at receptor complexes and adhesion sites (Elong Edimo et al., 2013; Thomas et al., 2017).

The N-terminal region of SHIP2 harbors the SH2 domain, a pleckstrin homology (PH)-related domain, and a RhoA-binding domain, all of which are embedded within an extended stretch predicted to contain intrinsically disordered segments (Jumper et al., 2021). While intrinsically disordered regions (IDRs) can facilitate weak, multivalent protein–protein interactions, structured domains can also mediate tight, stoichiometric dimerization (Chen et al., 2024). Our domain-mapping experiments show that a structured N-terminal fragment is both necessary and sufficient for homodimerization, whereas C-terminal fragments alone fail to interact with similar affinity. This contrasts with the structural logic underlying SHIP2 catalytic regulation, in which interdomain communication between the phosphatase domain and a proximal C2 domain is the primary determinant of catalytic efficiency (John et al., 2023; Le Coq et al., 2017). Together, these findings suggest that SHIP2 harbors at least two spatially distinct regulatory modules: the C2–phosphatase interface, which governs enzymatic turnover through an allosteric mechanism, and the N-terminal dimerization interface, which dictates its oligomeric form. We cannot, however, exclude the possibility that the N-terminal domain, while acting as the primary driver of homodimerization, simultaneously contacts a distal region within the C-terminal portion of the same or opposing protomer, thereby establishing a composite N-to-C dimerization interface. Such an architecture would be consistent with the relatively weak interaction signals observed in our experiments, as trans-domain contacts of this nature are often transient or of lower affinity than canonical symmetric interfaces.

In line with this hypothesis, deletion of the N-terminal dimerization domain or co-expression of ΔN-SHIP2 with wild-type SHIP2 results in only a moderate reduction in catalytic activity against PI (3,4,5) P3 in vitro, suggesting that the dimerization interface does not constitute an obligatory element for catalytic activity. This is mechanistically consistent with the known allosteric architecture of SHIP2, in which the C2 domain exerts a dominant stimulatory effect on the phosphatase domain via two independent allosteric pathways involving hydrophobic and polar interdomain contacts, without requiring any contribution from the N-terminal region (Le Coq et al., 2017). The mild reduction in activity we observe in the dimer-deficient and hetero-dimeric states is therefore most parsimoniously interpreted as a secondary, interdomain effect, possibly through steric perturbation of membrane docking or altered conformational dynamics of the full-length protein, rather than a direct loss of a catalytically essential element. This is reminiscent of the complex interdomain regulation in PTEN, where C-terminal tail phosphorylation and dimerization modulate membrane residence time and catalytic output through allosteric mechanisms that do not directly contact the active site (Masson and Williams, 2020). Altogether, our data suggest that while homodimerization may fine-tune SHIP2 activity, its primary function is likely to regulate protein–protein interactions and interactome selectivity rather than to switch catalytic function on or off.

One of the most important findings of this study is the profound and qualitative shift in the SHIP2 interactome that accompanies changes in its oligomeric state. Wild-type dimeric SHIP2 is preferentially associated with proteins involved in protein folding, cytoskeletal organization, and chaperone networks, while monomeric ΔN-SHIP2 is enriched in interactions with RNA-binding proteins (RBPs) and RNA metabolism factors, including G3BP1 and CAPRIN1. The shift from a predominantly signaling-competent interactome in the dimeric state to an RNA granule-associated interactome in the monomeric state raises the possibility that the dimerization equilibrium of SHIP2 operates as a molecular switch, redirecting its cellular activities in response to stress. This model further implies a functional antagonism between its two terminal domains: the N-terminal domain, by driving homodimerization, may favor engagement with the protein–protein interaction network, whereas the C-terminal domain may exert an opposing influence, orienting SHIP2 toward nucleic acid-associated functions, a partition that would only become apparent upon dissolution of the dimer. Another possibility is that deletion of the N-terminal domain of SHIP2 might expose interaction interface that are normally hidden in its native conformation, which in turn could engage in distinct protein-protein interaction. Interestingly, the interactome obtained from co-immunoprecipitation of wild-type SHIP2 in the presence of ΔN-SHIP2 closely resembles the ΔN-SHIP2 interactome, suggesting that the monomeric form may exert a dominant effect on interaction network remodeling when expressed alongside the dimeric form. This could reflect a competition for shared N-terminal interaction surfaces, or alternatively, a trans-acting sequestration mechanism in which ΔN-SHIP2 displaces dimeric SHIP2 from its canonical partners. Similar dominant-negative effects through disruption of the dimerization interface have been reported in other contexts, including the mutual inhibition of PTEN activity via domain-swap interactions (Masson and Williams, 2020).

The identification of G3BP1 as an SHIP2-interacting protein is particularly significant, given the central role of this protein in stress granule (SG) nucleation and assembly. G3BP1 and its paralog G3BP2 function as the essential nucleators of mammalian SGs; deletion of both G3BP paralogs almost completely abolishes SG formation under diverse stress conditions (Yang et al., 2020). Our domain-mapping experiments indicate that the interaction between SHIP2 and G3BP1 is mainly mediated by the N-terminal region of SHIP2, suggesting that SHIP2 dimerization and G3BP1 are mutually exclusive events at the N-terminal surface. In this context, a shift from the dimeric to the monomeric state of SHIP2 could increase the availability of its N-terminal domain for engagement by G3BP1, thereby linking SHIP2 conformational state to SG assembly. This model positions SHIP2 as a potential regulator of stress granule dynamics and provides a framework for future structural and mechanistic studies.

The functional experiments performed in MDA-MB-231 cells reveal that SHIP2 overexpression significantly increases the size of SGs formed in response to sodium arsenate, particularly for granules exceeding 3 µm in size, but does not affect their persistence after stress removal. Cells expressing phosphatase-dead SHIP2 show an intermediate phenotype relative to parental and wild-type SHIP2 cells, suggesting that both the adaptor function and the enzymatic activity of SHIP2 contribute to SG size and clearance kinetics. The partial contribution of phosphatase activity to this phenotype suggests that local PI (3,4,5) P3 hydrolysis at or near the condensate surface, potentially influencing membrane lipid composition or the recruitment of PI (3,4) P2-binding effectors, may tune SG dynamics, while the adaptor function of SHIP2 may independently influence SG composition or coalescence. Attempts to directly assess PI (3,4,5) P3 localization within stress granules using biosensors or antibody-based approaches proved technically challenging and did not yield sufficiently robust or reproducible data. Future studies will therefore be required to address how PI (3,4,5) P3 is dynamically regulated within or at the surface of SGs, and how the coordinated activities of lipid kinases and phosphatases contribute to this process.

Finally, our findings have important implications for therapeutic strategies targeting SHIP2. Indeed, several SHIP2 inhibitors have been developed and tested in preclinical models (Viernes et al., 2014); however, recent evidence indicates that some of these compounds exert SHIP2-independent effects, raising concerns about specificity (El Sayed et al., 2024). The finding that SHIP2 forms homodimers suggests that future inhibitor screening and design efforts should take its dimerization properties into account. Indeed, targeting dimer interfaces or stabilizing specific oligomeric states could provide a novel and more selective approach to modulating SHIP2 function. Such strategies may allow selective disruption of non-canonical SHIP2 functions while preserving or altering its catalytic activity.

In summary, our study identifies SHIP2 dimerization as a key determinant of its interactome and cellular functions, uncovering a previously unrecognized role for SHIP2 in stress granule regulation, and provides a conceptual framework for understanding how changes in SHIP2 expression and oligomerization may contribute to disease. These findings open new avenues for investigating the structural and functional plasticity of SHIP2 and related phosphoinositide phosphatases.

---

**Supplementary Figure 1:**
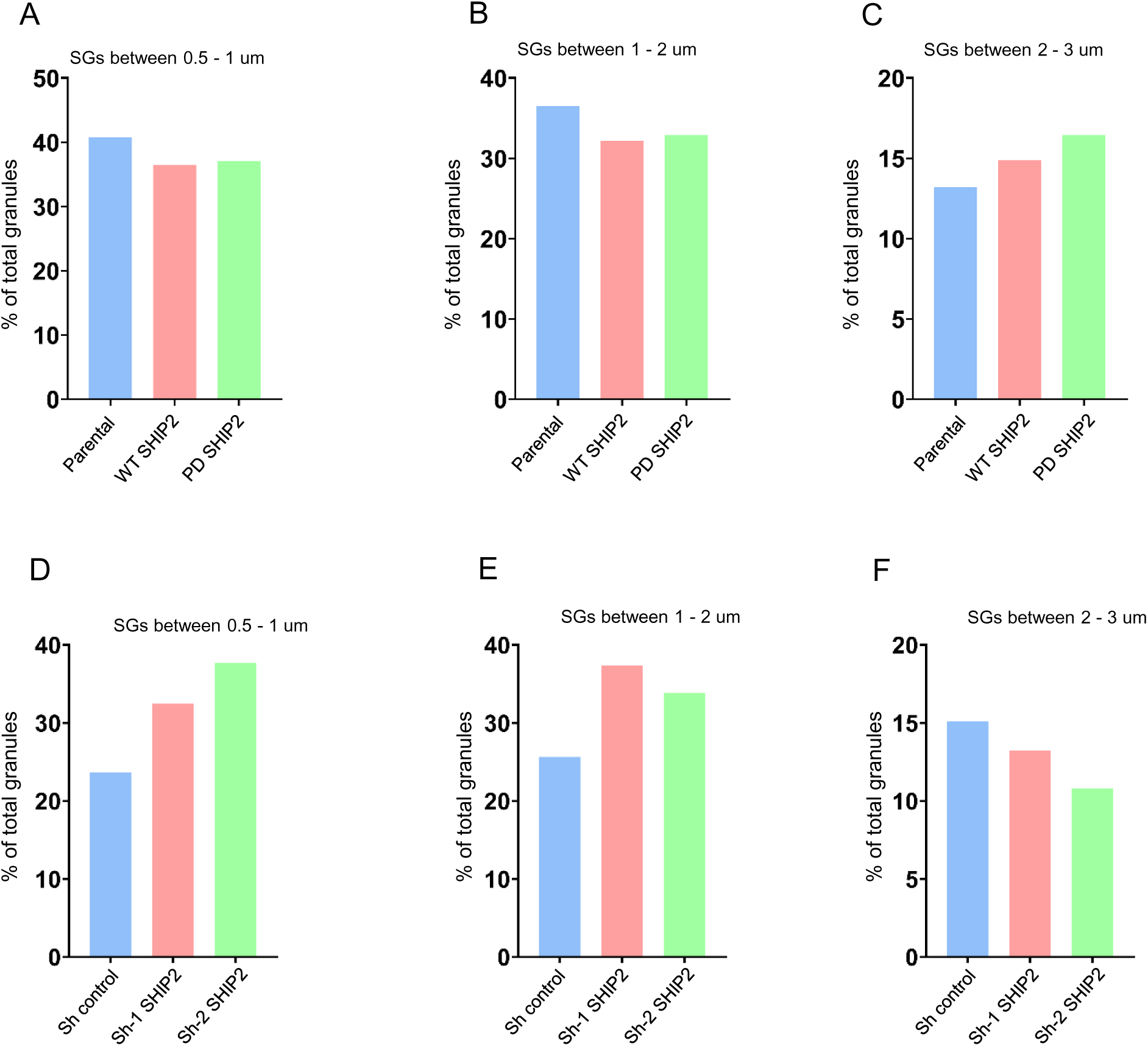
Quantification of SG size distribution after 1 hour of arsenate treatment Bar graphs representing the percentage of stress granules sizes between 0.5 and 3 um in cells Parental MDA-MB-231, and cells stably overexpressing wild-type SHIP2 (**A-C**) and cells expressing Sh control and SHIP2 knockdown using Sh1 and Sh2 **(D-F)**, the Y axis represents the percentage of the total granule pool, the X axis represents the cell line.

**Supplementary Figure 2:**
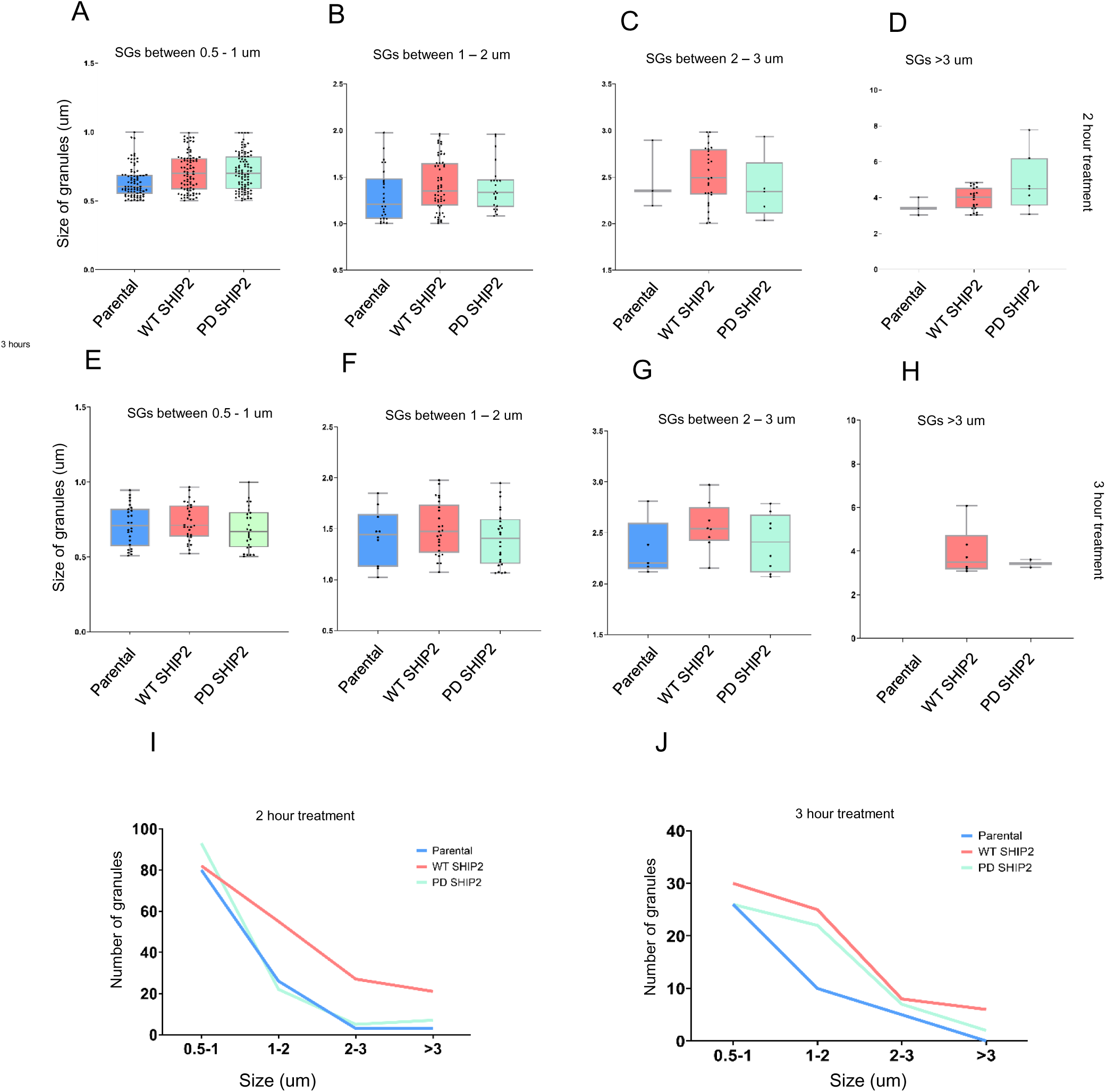
Quantification of SG size and distribution after 2 and 3 hours of arsenate treatment Box Blots representing the distribution of stress granules in cells Parental MDA-MB-231, and cells stably overexpressing wild-type SHIP2 after 2 (**A-D**) and 3 hours of treatment (**E-H**). the Y axis represents the size of stress granules, the X axis represents the cell line. (**I-J**) Line charts displaying the number of stress granules in cells Parental MDA-MB-231, and cells stably overexpressing wild-type SHIP2 after 2 (**I**) and 3 hours of treatment (**J**). the Y axis represents number of stress granules, the X axis represents the size.

## Supplementary tables

**Supplementary Table 1:** Quantitative interactome profiling of IP wild-type, heterodimeric, and dimer-deficient SHIP2 complexes.

Affinity Purification Mass Spectrometry (AP-MS) of immunoprecipitated Dimer Deficient (IP Flag Dimer Deficient), Heterodimer (IP His His+DD), and wild-type dimer SHIP2 (IP Flag His+Flag). Titles show Protein identification cards (Gene names, Protein IDs, Molecular weight, MS counts), intensity Based Absolute Quantification (iBAQ) values, and replicate intensity values (n=3 biological independent replicates per condition). Differential enrichment is reported as log2 fold-change (Log2FC) relative to the control, with statistical significance (-log10 p-value and q-value) from Student’s t-tests.

**Supplementary Table 2:** Gene Ontology Molecular Function enrichment analysis of IP wild-type, heterodimeric, and dimer-deficient SHIP2 complexes.

Columns indicate the specific GO term, gene overlap, significance metrics (P-value and Adjusted P-value computed via Fisher’s exact test).

**Supplementary Table 3:** Quantitative interactome profiling of wild-type, phosphatase dead, heterodimeric, and dimer-deficient SHIP2 complexes after arsenate treatment.

Affinity Purification Mass Spectrometry (AP-MS) of total cell lysate from Dimer Deficient (Dimer Deficient), Heterodimer (His+DD), Phosphatase dead (PD) and wild-type dimer (His+Flag). Titles show Protein identification (Gene names, Protein IDs, Molecular weight, MS/MS counts), intensity Based Absolute Quantification (iBAQ) values, and replicate intensity values (n=3 biologically independent replicates per condition). Differential enrichment is reported as log2 fold-change (Log2FC) relative to the control, with statistical significance (-log10 p-value and q-value) from Student’s t-tests.

**Supplementary Table 4:** Quantitative interactome profiling of endogenous SHIP2.

Affinity Purification Mass Spectrometry (AP-MS) of immunoprecipitated endogenous SHIP2. Titles show Protein identification (protein names, Protein IDs, Molecular weight, MS/MS counts), intensity Based Absolute Quantification (iBAQ) values, and replicate LFQ intensity values (n=3 biologically independent replicates per condition). Differential enrichment is reported as log2 fold-change (Log2FC) relative to the control, with statistical significance (-log10 p-value and q-value) from Student’s t-tests.

## Materials and methods

### Reagents

Dulbecco’s Modified Eagle Medium (DMEM) was purchased from CAPRICORN Scientific, cat. No. DMEM-HPA. Fetal Bovine Serum was purchased from Sigma Aldrich (#F7524). Penicillin/Streptomycin and 2.5% Trypsin (10X) were purchased from Gibco (#15140-122 and #15090-046). JetPRIME was purchased from Polyplus, #101000027.

### Cell culture

All cell lines were cultured in Dulbecco’s Modified Eagle Medium (DMEM) supplemented with 10% fetal bovine serum (FBS) and 100 U/mL penicillin/streptomycin. HEK293T and MDA-MB-231 cell lines were kindly provided by Dr. Anna Marusiak, MCF7 cell line by Dr. Karolina Szczepanowska, and the A375 cell line by Dr. Paweł Niewiadomski (Centre of New Technologies, University of Warsaw). All cell lines were maintained between passages 3 and 15. Cultures were grown in a humidified incubator at 37°C with 5% CO₂.

### Plasmid transfection

HEK293T cells were plated in 6-well dishes at 50 % confluency. Twenty-four hours later, the medium was replaced, and the cells were transfected with the appropriate plasmids using JetPRIME transfection reagent, following the manufacturer’s instructions. Briefly, the plasmids were resuspended in 200 µl of transfection buffer. The mixture was gently vortexed, Jetprime transfection reagent was added at a 1:2.5 ratio, vortexed again, and spun down, then incubated at room temperature for 10 min. The mixture was then added dropwise to each plate. Three to four hours later, the media was replaced by complete DMEM. The following plasmids were used: SHIP2-WT His/V5, SHIP2-ΔPS, SHIP2-ΔPR1, SHIP2-N1, SHIP2-N2, SHIP2-C1, SHIP2-C2, SHIP2-PD, SHIP2 WT Flag (a gift from Prof. Pavel Krejci, Department of Biology, Masaryk University, 62500 Brno, Czech Republic), and SHIP2 Δ-N was generated in-house. SHIP2 knockdown was performed using the plasmids described previously (El Sayed et al., 2024).

### Immuno-precipitation

HEK 293T cells were lysed on ice for 10 minutes in IP buffer containing 10% glycerol, 20 mM Tris (pH 7.4), 150 mM NaCl, 1 mM EDTA, 1.5 mM MgCl₂, and 0.5% IGEPAL, supplemented with 1 mM phenylmethylsulfonyl fluoride (PMSF) immediately before use. The lysates were centrifuged at 15,000 × g for 15 minutes at 4 °C. A fraction of the lysate (10%) was stored as input. Equal amounts of protein were transferred to new Eppendorf tubes. The corresponding antibodies were added, and samples were incubated on a rotating agitator for 3 hours at 4 °C. Roche Protein A-agarose beads (#11134515001) were washed three times with ice-cold PBS, followed by one wash with IP buffer. After, the beads were centrifuged at 300 × g for 3 minutes. Beads were dried by removing the maximum amount of buffer and re-suspended in an appropriate volume of lysis buffer before being added to the samples. Samples were incubated on a rotating shaker at 4 °C for an additional 2 hours. After incubation, samples were centrifuged, the lysate was removed, and the beads were washed three times with ice-cold PBS. After the last wash, the beads were resuspended in Laemmli buffer. Bead-bound proteins were denatured by boiling at 95 °C before loading onto SDS-PAGE gels. For native immunoprecipitation, cells from an 80% confluent 10 cm dish were lysed in IP buffer, and SHIP2 protein was immunoprecipitated using an anti-SHIP2 antibody.

### Western blot

Whole-cell lysates were prepared using a lysis buffer containing 50 mM Tris-HCl (pH 8.0), 150 mM NaCl, 0.5% sodium deoxycholate, 1.0% NP-40, 0.1% SDS, and 1 mM phenylmethylsulfonyl fluoride (PMSF). Protein extracts were incubated on ice for 10 minutes and then centrifuged at 20,000 × g for 12 minutes at 4 °C. Protein concentration was determined using Pierce™ BCA Protein Assay Kit (Thermo Scientific™, #23225), following the manufacturer’s instructions. For protein separation, sodium dodecyl sulfate-polyacrylamide gel electrophoresis (SDS-PAGE) was performed, and proteins were subsequently transferred onto either nitrocellulose or polyvinylidene fluoride (PVDF) membranes. Membranes were blocked at room temperature using Tris-buffered saline (TBS) containing 0.05% Tween-20 (TBS-T) and 5% non-fat milk. Membranes were incubated with primary antibodies overnight at 4 °C with continuous rotation. The following primary antibodies were used: phospho-AKT1 (Ser473) (#700256), ERK1/ERK2 (#13-6200), Phospho-ERK 1/2 (#14-9109-82), SHIP2 (#MA5-14844), AKT Pan (#MA5-14916), 6x His tag (#MA1-135), Anti-FLAG (#F1804-200UG), P70 S6 kinase (#MA5-15141), Phospho- P70 S6 kinase (#710095), EIF2s1 (#AHO0802), G3BP1 (#PA5-29455), Anti-V5 tag (#MCA1360GA). Antibody dilutions were prepared according to the manufacturer’s recommendations. Chemiluminescent signals were detected using the Amersham ImageQuant 800 system.

### Phosphatase assay

SHIP2 phosphatase activity was measured using phosphatase assay buffer consisting of 25 mM Tris-HCl (pH 7.4), 140 mM NaCl, 2.7 mM KCl, and 5 mM DTT, with the Malachite Green Phosphate Assay Kit (Sigma-Aldrich, MAK307) according to the manufacturer’s instructions. A phosphate standard curve was prepared by serial dilution of the provided 1 mM phosphate standard in ultrapure water, yielding concentrations ranging from 0 to 40 µM. The working reagent was prepared by mixing Reagent A and Reagent B at a 100:1 ratio and equilibrated to room temperature prior to use. 80 uL of each sample or standard was transferred in duplicate to a clear flat-bottom 96-well plate, followed by the addition of 20 µL of working reagent. After gentle mixing, the plate was incubated at room temperature for 30 minutes to allow color development. Absorbance was measured at 620 nm using a spectrophotometric microplate reader, and phosphate concentrations in each sample were calculated from the standard curve.

### Cloning and Mutagenesis

An N-terminal deficient SHIP2 construct (ΔN-SHIP2, Δ21-427aa) was generated using the Q5® Site-Directed Mutagenesis Kit (New England Biolabs NEB # E0554S) according to the manufacturer’s protocol. SHIP2-WT Flag plasmid was amplified using the following forward: ATAGGCACCTGGAAC and reverse: GGAGGGGGCCTGGCT primers designed using the NEBaseChanger Tool, https://nebasechanger.neb.com/.

### Mass Spectrometry Sample Preparation

Protein LoBind Eppendorf tubes were used to collect cell pellets. Pellets were washed twice with PBS and dried by removing PBS. The cell lysate was prepared by adding Trifluoroacetic Acid (TFA) in a 1:4 pellet/TFA ratio and incubated for 10 minutes at room temperature with occasional vortexing. Proteins were precipitated using a neutralization Buffer (2M TrisBase) in a 10:1 ratio (neutralization buffer/TFA). Next, a 10X Reduction/Alkylation Solution containing 100 mM Tris(2-carboxyethyl) phosphine and 400 mM 2-chloroacetamide was added at a 1.1:1 reduction/alkylation/TFA ratio. Samples were incubated at 95°C for 5 minutes. Protein concentration was determined using turbidity measurement (McFarland). The protein content of each sample was adjusted to 1 µg/µL using a Sample Dilution Buffer composed of 10:1 (v/v) 2M Tris Base/TFA. Samples were further diluted 1:5 in H2O; trypsin was added, and the mixture was incubated at 37°C under constant vortexing at 950 rpm overnight in a ThermoMixer. After digestion, samples were centrifuged at max speed for 2 minutes and transferred to new protein LoBind tubes; TFA was added to a final concentration of 2%, and the pH was measured to be ∼2.

### IP Sample preparation

Following immunoprecipitation (IP), proteins bound to the beads were digested overnight with trypsin using the same protocol (on-bead digestion). The samples were then centrifuged at maximum speed for 2 minutes, and the resulting supernatant was collected and transferred to new protein LoBind tubes. Trifluoroacetic acid (TFA) was added to a final concentration of 2%.

### Tandem Mass Tag (TMT) Labelling and Fractionation

Columns for labeling were prepared on Stop And Go Extraction STAGE tips packed with three layers of C18 mesh. TMT labels were prepared by dissolving them in 100% acetonitrile (ACN) and adding 200 µl 50 mM Na2PO4/NaH2PO4, pH 8.0, to each label. STAGE tips were first conditioned with 150 µl methanol and centrifuged at 1200 g for 2 mins at room temperature; next, the resin was washed with 100 µl of 50% acetonitrile in 0.1% Formic Acid (FA). The resin was then equilibrated twice by adding 150 µl of 0.1% FA and centrifuging. 10 µg of digested peptides were loaded, then washed twice with 150 µl of 0.1% FA. 200 µl of the corresponding TMT label was added, and the mixture was centrifuged at 300g for 15 minutes or until the TMT solution passed through. Samples were then washed three times with 150 µl of 0.1% FA. STAGE tips were later transferred to new protein LoBind tubes, and the samples were eluted by adding 60 µl of 60% ACN in 0.1% FA. Then, 55 µl of each sample was mixed in a new Eppendorf tube and evaporated using a SpeedVac at 40°C overnight. Samples were then fractionated using Pierce High pH Reversed-Phase Peptide Fractionation Kit (ref#) according to the manufacturer’s instructions.

### TMT-based differential analysis of protein levels in total cell extracts

Mass spectrometry analysis yielded a list of identified peptides for each condition. Raw intensity values were normalized to the control condition, consisting of non-treated cells transiently transfected with an empty pcDNA-FLAG vector. Enrichment thresholds were defined as a fold change of ≥ 1 (for both enrichment and depletion) and a −log₁₀ p-value of ≥ 1.3.

### Immunofluorescence

Cells were plated on glass coverslips (treated or not with 0.001 % Poly-L-lysine (Sigma Aldrich, #P4832) for 1 hour) and left to adhere overnight in 12-well plates (100,000 cells/well). Adherent cells were then transfected with relevant plasmids. Transfection was carried out using JetPRIME transfection reagent (Polyplus, #101000027) according to the manufacturer’s instructions. 4 hours later, the media was changed. Cells were then fixed with 4% freshly prepared paraformaldehyde (Carl Roth). The coverslips were then washed three times with PBS (10 minutes each wash) and were then incubated for 1 hour at room temp in Blocking buffer (5% Goat serum, 0.3% triton X100), then washed with PBS and incubated with the corresponding antibodies (diluted according to manufacturer’s instructions) diluted in antibody-dilution buffer (1% BSA, 0.3% TritonX) overnight at 4 C. the coverslips were then washed three times with PBS before incubating for 1 hour at room temp with the respective secondary antibodies conjugated to a fluorofore (Alexafluor 488/568 cat # A-11008 and # A-11011 respectively). The slides were later washed at room temp with PBS three times and once with distilled water, then mounted on glass slides using ProLong™ Diamond Antifade Mountant (Thermo Fisher, cat #P36970) and left to dry. The images were captured using a Zeiss LSM 910 confocal microscope and processed with ImageJ (National Institutes of Health, Bethesda, MD, USA).

### Confocal Image Analysis

Confocal images were acquired using a Zeiss LSM 910 confocal microscope. The images were exported from Zen to TIFF format.

### Image Analysis of Stress Granule Morphology

Fluorescence micrographs were analyzed using Fiji (ImageJ, NIH). For each image, a region of interest (ROI) was manually defined to restrict analysis to the relevant area. The image was duplicated to preserve the original, and background fluorescence was subtracted using a rolling-ball algorithm with a 35-pixel radius. Pixels outside the ROI were cleared prior to further processing. Images were converted to 16-bit format, and a spatial calibration of 187 pixels per 10 µm was applied. Thresholding was performed using the Otsu algorithm on the dark background to reliably capture intensely fluorescent granules while excluding dim or diffuse signal. Thresholded images were converted to binary masks with a black background convention. To separate close granules, a watershed segmentation algorithm was applied to the binary mask. The resulting watershed image was saved as a TIFF file. Particle analysis was then performed on the segmented mask using the “Analyze Particles” function, with a minimum granule size cutoff of 0.04 µm² and no upper size limit. Particles touching the image border were excluded from quantification. For each detected granule, area, mean fluorescence intensity, and integrated density were recorded. Results were exported as CSV files for downstream statistical analysis, and detected granules were added to the ROI Manager for visual verification.

### Statistics

We used GraphPad Prism (http://www.graphpad.com) for all statistical analyses. Each experiment included at least three independent biological replicates. To compare two groups, we applied an unpaired, two-tailed t-test. For evaluating three or more groups, we utilized a one-way ANOVA followed by Dunnett’s post hoc test. We defined statistical significance as P < 0.05.

## DECLARATIONS

### Declaration of generative AI and AI-assisted technologies in the manuscript preparation process

The AI tools Grammarly and ChatGPT were used in the final stage of manuscript preparation for English corrections and clarifications. After using these tools, the author(s) reviewed and edited the content as needed.

### Ethics approval and consent to participate

not applicable.

### Consent for publication

not applicable.

### Availability of data and materials

The datasets used and/or analyzed during the current study are available from the corresponding author on reasonable request.

### Authors’ contributions

Conceptualization, A.A, Methodology; A.A and A.E.S, Investigation, A.A and A.E.S.; original draft preparation, A.E.S and AA; Writing review and editing, A.A and A.E.S; Funding acquisition, A.A.

## Acknowledgements

Special thanks to Dr. Remigiusz Serwa and Dr. Dorota Stadnik for help with sample preparation, LC-MS/MS measurements, and raw data processing. Special thanks to Prof. Pavel Krejci (Department of Biology, Masaryk University, 62500 Brno, Czech Republic) for providing the SHIP2 plasmid constructs.

